# Improving the study of RNA dynamics through advances in RNA-seq with metabolic labeling and nucleotide-recoding chemistry

**DOI:** 10.1101/2023.05.24.542133

**Authors:** Joshua T. Zimmer, Isaac W. Vock, Jeremy A. Schofield, Lea Kiefer, Michelle H. Moon, Matthew D. Simon

## Abstract

RNA metabolic labeling using 4-thiouridine (s^4^U) captures the dynamics of RNA synthesis and decay. The power of this approach is dependent on appropriate quantification of labeled and unlabeled sequencing reads, which can be compromised by the apparent loss of s^4^U-labeled reads in a process we refer to as dropout. Here we show that s^4^U-containing transcripts can be selectively lost when RNA samples are handled under sub-optimal conditions, but that this loss can be minimized using an optimized protocol. We demonstrate a second cause of dropout in nucleotide recoding and RNA sequencing (NR-seq) experiments that is computational and downstream of library preparation. NR-seq experiments involve chemically converting s^4^U from a uridine analog to a cytidine analog and using the apparent T-to-C mutations to identify the populations of newly synthesized RNA. We show that high levels of T-to-C mutations can prevent read alignment with some computational pipelines, but that this bias can be overcome using improved alignment pipelines. Importantly, kinetic parameter estimates are affected by dropout independent of the NR chemistry employed, and all chemistries are practically indistinguishable in bulk, short-read RNA-seq experiments. Dropout is an avoidable problem that can be identified by including unlabeled controls, and mitigated through improved sample handing and read alignment that together improve the robustness and reproducibility of NR-seq experiments.

## Introduction

RNA-sequencing (RNA-seq) is a standard experiment to characterize the expression profile of cells and quantify changes in expression. Unfortunately, traditional RNA-seq experiments provide minimal information about changes in mRNA stabilities or synthesis rates. RNA metabolic labeling can be used to study the dynamics of RNA populations. In these experiments, cells are treated for a fixed period of time with noncannonical nucleosides such as s^4^U, which are incorporated into RNA and identified either via enrichment-based approaches (Cleary et al. 2005; Dölken et al. 2008; Duffy et al. 2015; Duffy et al. 2018; Schwalb et al. 2016) or through enrichment-free RNA sequencing-based approaches (Herzog et al. 2017; Riml et al. 2017; Schofield et al. 2018; Kiefer et al. 2018; Chen et al. 2020; Gasser et al. 2020). These enrichment-free methods (e.g., SLAM-seq, TUC-seq, TimeLapse-seq) often use 4-thiouridine (s^4^U) and/or 6-thioguanosine (s^6^G) to metabolically label newly synthesized RNA for an extended period of time (typically 1-3 h), as had been developed previously for methods based on enrichment (Neymotin et al. 2014). Upon purifying RNA, the s^4^U is chemically converted from a uridine analog to a nucleotide that reverse transcriptases read as a cytidine (and analogously s^6^G to adenine), so we refer to this family of experiments as nucleotide-recoding RNA-seq (NR-seq) technologies. In TimeLapse, s^4^U is oxidized with a reagent such as sodium periodate (NaIO_4_) to a reactive intermediate and undergoes nucleophilic attack with an amine such as 2,2,2-trifluoroethylamine (TFEA). TUC chemistry is very similar but employs osmium tetroxide (OsO_4_) as the oxidant and ammonia (NH_3_) as the amine. Both TimeLapse and TUC chemistry completely recode the H-bonding pattern of s^4^U from that of a uridine analog to that of a cytidine analog. In SLAM chemistry, s^4^U is alkylated with iodoacetamide (IAA) which disrupts the H-bonding pattern. In all these methods, when the cDNA from chemically treated RNA is sequenced and aligned to the appropriate genome, sites of label incorporation manifest as an apparent mutation. T-to-C (or G-to-A) mutations are then used to infer the population of newly synthesized RNA and the proportion of newly synthesized RNA can be used to estimate the half-life of the transcript (Schofield et al. 2018; Jürges et al. 2018).

While NR-seq and other metabolic labeling approaches have proven powerful, we and others have noted that labeled reads appear underrepresented in some samples compared with unlabeled controls (Schofield et al. 2018; Berg et al. 2023), a phenomena we refer to as dropout. Here we investigate dropout across a number of potential origins revealing dropout can be driven by specific loss of s^4^U-labeled transcripts at two distinct steps: (1) handling of the RNA samples and (2) incomplete alignment during analysis. We demonstrate both sources of dropout can be addressed to minimize bias in NR-seq experiments. Importantly, this dropout is observed independently of any biological effects of s^4^U or which NR-seq chemistry is used. We demonstrate TimeLapse-seq, SLAM-seq and TUC-seq recoding efficiencies are similar, do not substantially influence dropout, and provide strongly concordant estimates of RNA degradation rate constants. Dropout during handling of the RNA samples occurs because s^4^U-containing RNA is lost more than unlabeled RNA during handling, especially when exposed to surfaces of cell culture dishes and untreated test tubes. This problem can be addressed by following best practices when handling RNA samples, namely avoiding cell lysis in cell culture dishes and using test tubes designed to minimize nucleotide surface adherence. We find that computational dropout in NR-seq data occurs because standard alignment software assume all mismatches should penalize alignment scores. These penalties force highly mutated reads to drop below filter cutoffs and artificially lower the proportion of mutation-containing reads in processed data. Computational dropout can be addressed using a 3-nucleotide (3-nt) alignment strategy, which has already been demonstrated to be important when aligning very short reads containing T-to-C chemically induced mutations (Zimmer et al. 2021). These two solutions increase the yield of reads from label-containing transcripts and improve robustness of NR-seq protocols. The considerations presented here are important for producing high-quality NR-seq datasets and take full advantage of the kinetic information provided by these methods.

## Results

### Handling s^4^U-labeled RNA to avoid dropout

The nucleoside s^4^U is a common reagent to metabolically label RNA, and many methods have been developed to take advantage of its chemical properties (Cleary et al. 2005; Kenzelmann et al. 2007; Dölken et al. 2008; Neymotin et al. 2014; Herzog et al. 2017; Riml et al. 2017; Schofield et al. 2018; Duffy et al. 2019). Despite this, we are not aware of any studies characterizing the behavior of s^4^U-containing RNA compared to unlabeled RNA. We were motivated to investigate the relative behavior of labeled and unlabeled RNA due to observation that fast turnover transcripts (i.e. those expected to have high s^4^U-incorporation rates and high numbers of T-to-C mutation-containing reads in NR-seq data) are under-represented in s^4^U-labeled samples compared to unlabeled samples (Figure 1A). While incomplete extraction of RNA including nascent and chromatin-bound RNA is well documented (Chujo et al. 2017), incomplete extraction would not explain differences between labeled and unlabeled samples. We noted that this trend was pronounced in samples with high s^4^U incorporation rates and long s^4^U treatment times in which reads associated with fast-turnover (i.e., highly labeled) transcripts such as *MYC* or *MIR17HG* were under represented relative to reads associated with stable (i.e., minimally labeled) transcripts such as *GAPDH* or *RPL5* (from Schofield et al. 2018, Figure 1A), and the loss is observed even when the RNA is not treated with any NR-seq chemistry. We analyzed four publicly available NR-seq datasets with matched unlabeled samples using a range of chemistries and found this phenomena is widespread (Narain et al. 2021; Fisher et al. 2022; Richters et al. 2021; Zuckerman et al. 2020). For three of the four datasets, we observed the same s^4^U-dependent read loss over genes with high proportions of T-to-C mutation-containing reads, and the effect was more pronounced with longer labeling times (Figure 1B,C). We hypothesized that apparent loss of reads from s^4^U-labeled RNA is a common and unaddressed problem that could be caused by dropout of s^4^U during RNA sample handling or computational dropout during alignment of NR-seq reads.

**Figure 1:**
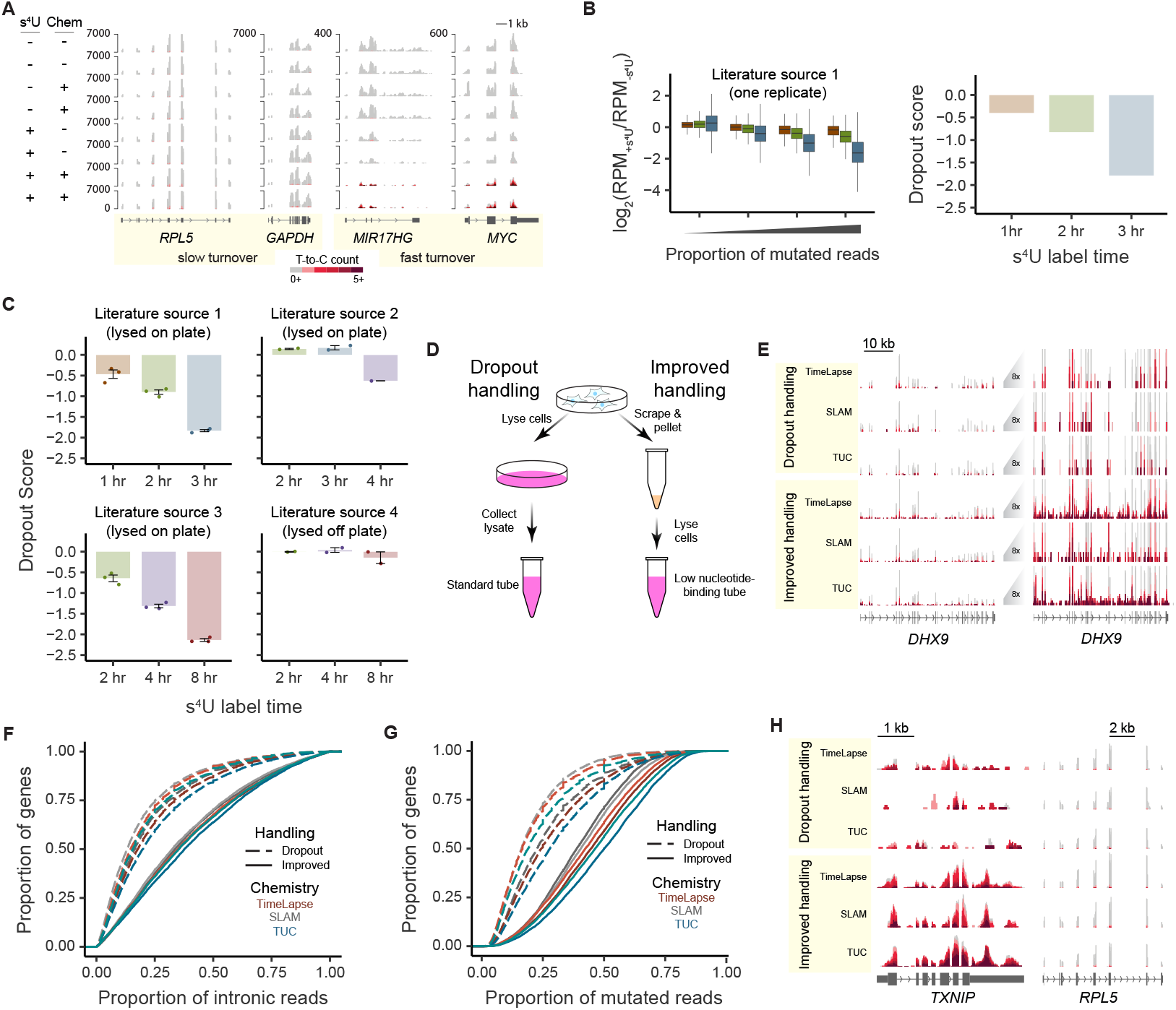
s^4^U-labeled RNA is specifically lost during handling. (A) Example NR-seq tracks from an experiment with high levels of dropout from K562 cells treated with s^4^U for 4h from Schofield et al. 2018. Reads aligning to transcripts with fast turnover (*MYC, MIR17HG*) are relatively under represented in the s^4^U treated samples whereas slow turnover transcripts (*GAPDH, RPL5*) are not. Tracks are colored by increasing numbers of apparent T-to-C mutations. (B) To quantify dropout, genes were grouped by the proportion of reads containing a mutation in NR-seq data and compared to the log_2_ fold difference of reads per million in samples labeled with s^4^U relative to those which were not. The dropout score is defined as the difference in mean log_2_ fold differences for transcripts with the highest and lowest proportion of mutated reads. (C) Assessing dropout score in four publicly available datasets. (D) Schematic of the dropout-vulnerable RNA isolation approach to one that improves s^4^U-labeled RNA yield. (E) Example NR-seq tracks of coverage over a highly intronic gene collected with both handling approaches and treated with TimeLapse, SLAM, or TUC chemistry. (F and G) Cumulative distribution plots of the proportion of reads aligning to an intronic region (F) or proportion of reads containing a T-to-C mutation (G) for all genes in NR-seq data. (H) Example NR-seq tracks of coverage over the gene of a high-turnover (TXNIP) and slow-turnover (RPL5) transcript.

While considering the apparent loss of s^4^U-labeled RNA, we noted samples with significant dropout (literature sources 1, 2, and 3 (Narain et al. 2021; Fisher et al. 2022; Richters et al. 2021)) and some of our earlier NR-seq datasets were collected using a protocol in which cells were lysed directly in cell culture dishes. This observation lead us to speculate that s^4^U-labeled RNA is more vulnerable to adhering to cell culture dishes and untreated plastics and that RNA sample handling could contribute significantly to dropout. To address the possible loss of labeled RNA during extractions, we developed a new handling protocol to improve the recovery of s^4^U-containing transcripts (Figure 1D). We avoided directly lysing cells in cell culture dishes because this method is susceptible to specific dropout of s^4^U-containing RNA, apparent in sequencing tracks for *DHX9* (Figure 1E). We considered an updated protocol to handle s^4^U-containing RNA that involves scraping cells from the cell culture dish and lysing in low nucleotide-binding sample tubes (Figure 1D).

To assess the extent to which our alternative protocol limits dropout, we treated adherent cells with s^4^U for 2 hours and collected RNA with both protocols. We performed chemistry developed for TimeLapse-seq, SLAM-seq, or TUC-seq on s^4^U-labelled RNA from the same samples to further test if the effect of the alternative protocol is chemistry-dependent. Upon examining the data, we found that NR-seq data correlate well with other data collected with the same handling but not with data collected with the other conditions, both in terms of total reads and T-to-C mutation-containing reads (Figure S1A,B). In addition, highly-labeled RNA and nascent transcripts containing introns are visibly depleted from samples collected under dropout conditions when compared to data collected with the improved handling (Figure 1E). To quantify the loss of highly-labeled transcripts we calculated the proportion of intronic reads aligning to each gene as we expect these to be essentially 100% labeled due to their short half-lives relative to the 2 hour labeling period. This analysis demonstrated that nascent RNA is globally underrepresented in the dropout dataset compared to the improved handling dataset independent of the conversion chemistry (Figure 1F). To test the effect on all s^4^U-containing RNA, we calculated the proportion of all reads aligning to each gene which contain a T-to-C mutation (Figure 1G). We found that sequencing data from improved handling conditions contain a higher proportion of mutation-containing reads, suggesting that these conditions significantly reduce dropout compared to standard protocols. The improvement was also evident in sequencing tracks by comparing fast turnover (*TXNIP*) and slow turnover (*RPL5*) transcripts. We found that the relative coverage over fast-turnover transcripts is strikingly increased with the improved handling protocol and slow-turnover transcripts are relatively unaffected (Figure 1H). This shows that specific dropout of s^4^U-labeled RNA biases NR-seq data, leading to an overrepresentation of previously existing RNA. Ultimately, this will shift estimates of transcript half-lives and turnover kinetics, the major advantage of employing NR-seq approaches. Our improvements to NR-seq protocols reduce bias in estimates for kinetic parameters and show that dropout is an issue with s^4^U, not specific to the NR-seq chemistry used.

### 3-nt alignment reduces computational dropout in NR-seq data

We considered the possibility that RNA sample handling might not be the only source of dropout and that some dropout might be computational, occurring during alignment. Most traditional sequencing aligners incur an alignment score penalty when a mismatch is identified within a read. Therefore, NR-induced mutations inadvertently decrease the chance a read will be successfully aligned. Most alignment softwares are customizable to allow for mismatch penalty minimization, but this simultaneously reduces the penalty for mismatches not caused by chemical conversion of the nucleotide analog. Furthermore, we previously demonstrated that a single chemically induced T-to-C mutation in small RNAs is sufficient to cause misalignment (Zimmer et al. 2021). We reasoned that the same effect may also be observed in longer sequencing reads.

Recently, HISAT-3N was developed as a 3-nt version of the splice-aware aligner HISAT2 which is commonly used to align RNA-seq data (Kim et al. 2015; Zhang et al. 2021). 3-nt alignment approaches intentionally convert all instances of one nucleotide to another within the sequencing reads and reference genome. In the case of NR-seq data using T-to-C chemistry, all T’s are converted to C’s, thereby masking chemically induced T-to-C conversions. HISAT-3N is the first 3-nt aligner specifically developed for NR-seq data, and while the authors previously validated its accuracy and efficiency, the recovery of mutation-containing reads in NR-seq data was not tested. Therefore, we aligned our improved handling NR-seq data using HISAT-3N and two popular 4-nt aligners: HISAT2 and STAR (Dobin et al. 2013). We explored relaxing the mismatch stringency of the 4-nt aligners, testing both moderate and extreme amounts of mismatch leniency (Figure 2A). These datasets allowed us to compare the ability of 3-nt and 4-nt aligners to recover s^4^U recoding induced mutations.

**Figure 2:**
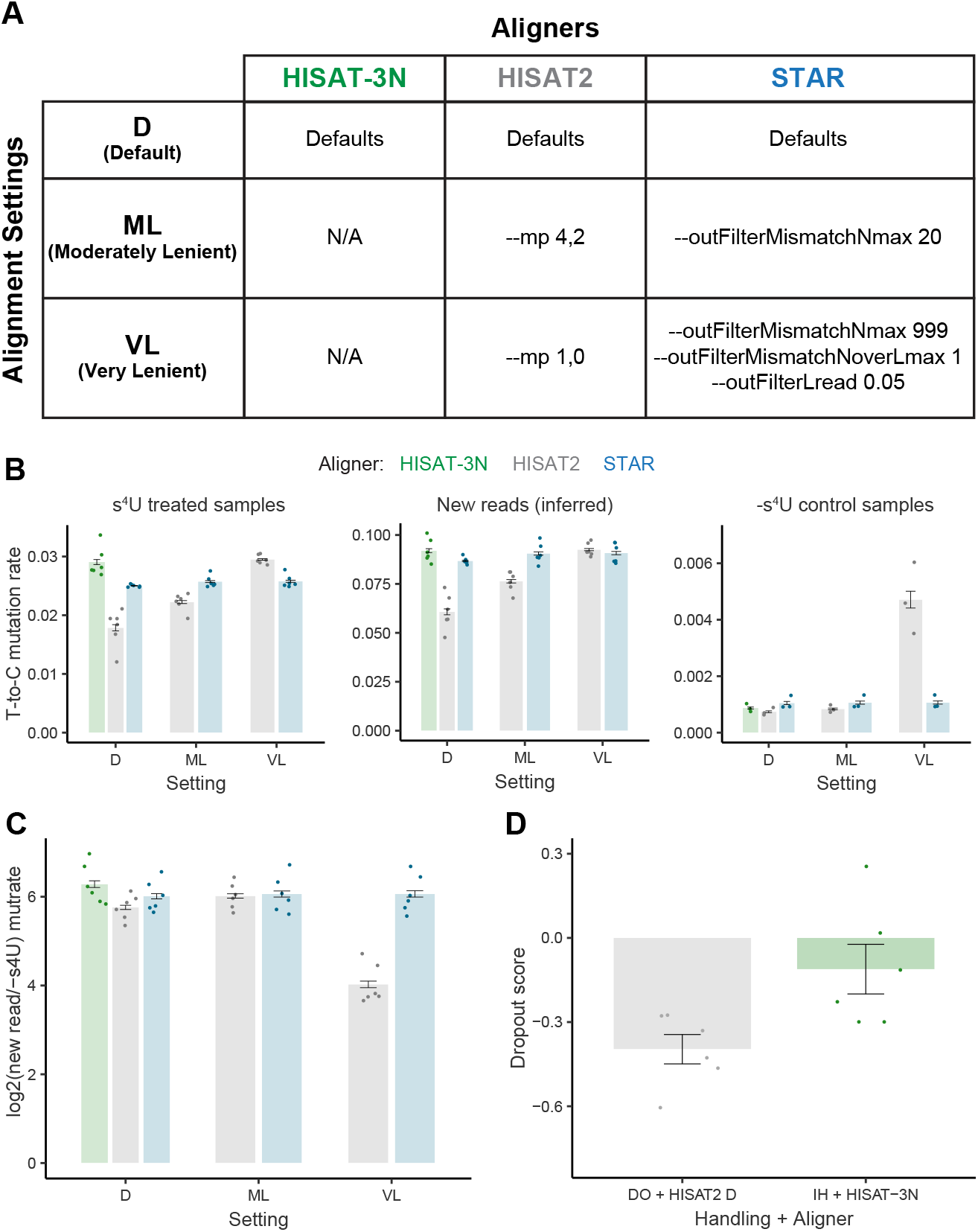
3-nt alignment recovers s^4^U recoding induced mutations. (A) The set of aligners and alignment settings used in this study. (B) Left: Sample-wide average mutation rates for all reads from s^4^U treated samples. Mutation rates are normalized to the average for each chemistry to remove batch and chemistry-specific effects. Middle: Mutation rate in new reads, inferred with mixture modeling and normalized as in the left plot. Right: Sample-wide average mutation rates for all reads from unlabeled samples. In all plots, points represent data for 2 (+s^4^U) or 1 (-s^4^U) replicate of each of the three NR chemistries considered. Error bars are +/- 1 standard error. (C) Difference between the new and old read mutation rates. (D) Impact of improved handling and alignment on dropout score (as defined in Figure 1B).

We assessed the T-to-C mutation rates in s^4^U treated samples. We found that when using default alignment parameters, HISAT-3N yielded higher mutation rates than either HISAT2 or STAR (Figure 2B left). We used mixture modeling to infer the mutation rate of reads from new RNA (Vock et al. 2023). New read mutation rates were also higher with HISAT-3N than default HISAT2 and STAR (Figure 2B middle). These differences were driven by HISAT-3N’s successful alignment of high T-to-C mutation content reads (Figure S2). Increasing the number of allowed mismatches in STAR did not significantly impact recovery of mutation containing reads. Relaxing HISAT2’s mismatch penalization led to increases in both sample-wide and new read mutation rates. HISAT-3N is thus able to recover more high mutation content reads than 4-nt aligners without globally reducing mismatch penalization.

We hypothesized that while HISAT-3N was likely recovering true s^4^U conversion induced mutations, HISAT2 with greater mismatch leniency was likely recovering a mixture of true and false positive mutations. To test this hypothesis, we compared mutation rates in samples not treated with s^4^U (i.e., -s^4^U control samples). In support of our hypothesis, while HISAT-3N’s background mutation rate was not significantly higher than with default 4-nt alignment, HISAT2 with extreme mismatch leniency saw a large increase in the sample-wide and transcript-specific background mutation rates (Figure 2B right and S2). Higher new read mutation rates can make identifying bona fide new reads easier, but only if the background mutation rates stay relatively low. To assess the extent of separation between new and old read mutational distributions, we calculated the log_2_-fold difference in the new and old read mutation rates for each aligner/setting combination. We found that HISAT-3N outperformed all 4-nt aligners by this metric, regardless of the extent of mismatch leniency in the latter (Figure 2C). Finally, we found that HISAT-3N’s use in conjunction with improved RNA handling drastically reduced dropout (Figure 2D) addressing both challenges for NR-seq data such as shown in Figure 1. Therefore, we conclude that HISAT-3N is the splice-aware aligner most appropriate for NR-seq data.

Similarly to dropout caused by RNA sample handling, the biases introduced by computational dropout will artificially lower the total proportion of mutation-containing reads, and more strongly affect reads derived from high-turnover transcripts (Berg et al. 2023). In addition, computational dropout is not specific to any NR-seq chemistry as it only depends on the presence of NR-induced mutations. Therefore, we recommend that all NR-seq data be aligned using a 3-nt approach using HISAT-3N or a similar aligner that does not penalize chemically-recoded bases (Sedlazeck et al. 2013; Neumann et al. 2019). Alternatively, a strategy to recover reads rendered difficult to map due to induced mutations was recently implemented in the NR-seq analysis software GRAND-SLAM (Berg et al. 2023), though HISAT-3N has the advantage of being fully open-source. Finally, other software such as Bismark in combination with Bowtie 2 can be used when splicing information is not required (Krueger et al. 2011; Langmead et al. 2012; Zimmer et al. 2021).

### Exploration of TimeLapse chemistry with s^4^U

Improvements in handling s^4^U-containing RNA and NR-seq data analysis will allow us to be more confident that we are observing all chemically converted s^4^U incorporation sites. To take advantage of these improvements we explored ways to increase the mutational content of reads from s^4^U labeled RNA by determining optimal TimeLapse reaction conditions with biological samples. Standard TimeLapse chemistry is performed under slightly acidic conditions (pH 5.2) to avoid basic conditions that would promote RNA hydrolysis. Sodium periodate (NaIO_4_) is used as the oxidant and 2,2,2-trifluorethylamine (TFEA) as the nucleophilic amine because our *in vitro* restriction endonuclease assay and NMR experiments suggested this is the most efficient oxidant-amine combination under conditions which do not promote RNA hydrolysis (Schofield et al. 2018). In addition, NaIO_4_ is a commonly used oxidant in RNA molecular biology and the low pKa leads to TFEA remaining mostly deprotonated in slightly acidic to neutral buffers.

First, we performed TimeLapse-seq in duplicate with labeled RNA purified using improved handling. We varied the reaction pH to compare standard conditions (pH 5.2) to a near-neutral buffer (pH 7.4). We examined the mutational content of all reads aligning to intronic regions as they are expected to be almost entirely labeled after a two hour treatment. The total proportion of mutation-containing intronic reads and average mutations per U were higher under more acidic conditions, demonstrating that the reaction is more efficient under slightly acidic conditions despite a higher fraction of protonated TFEA (Figures 3A & S3A).

**Figure 3:**
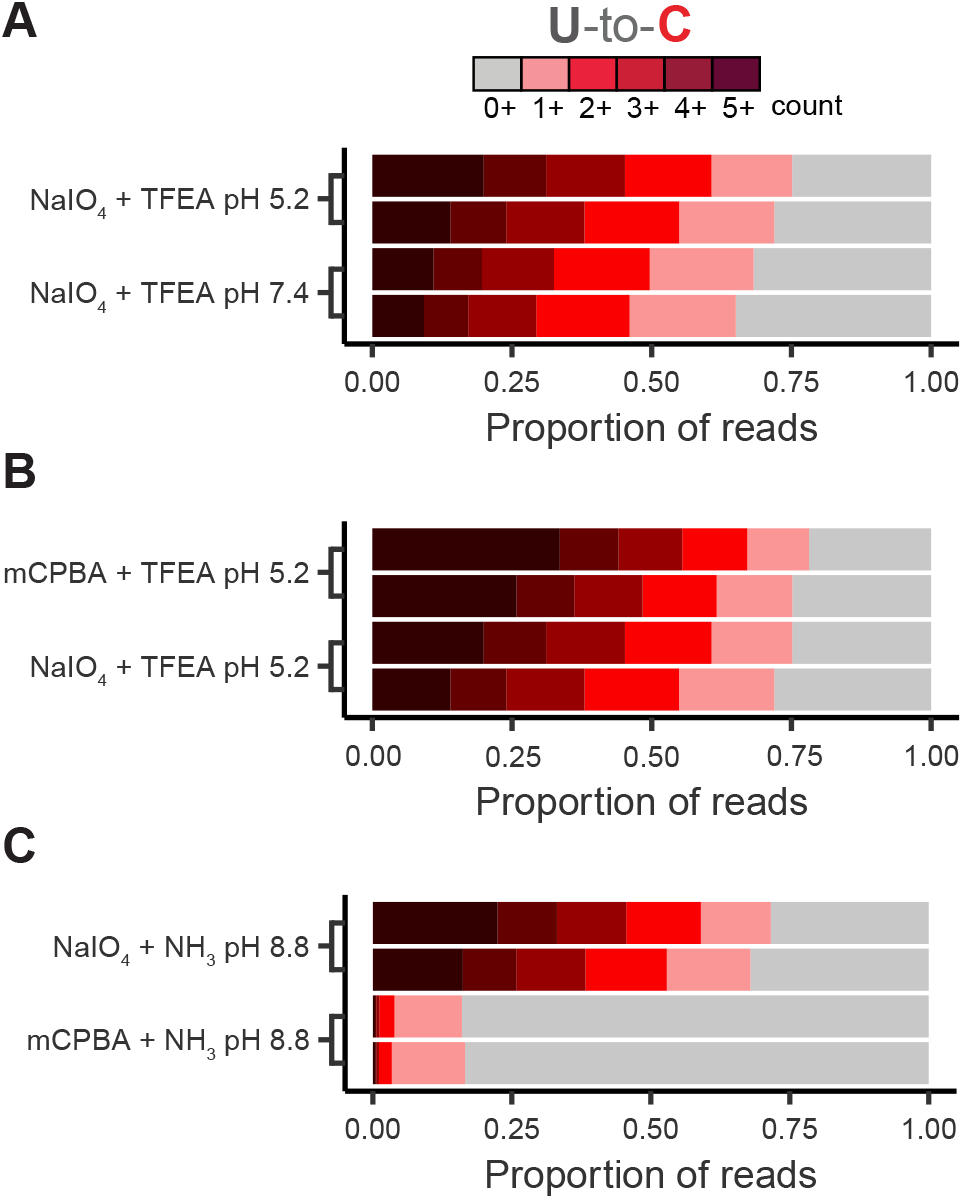
Buffer conditions and reagents affect efficiency of TimeLapse chemistry. (A-C) The proportion of intron-aligning reads which contain T-to-C mutations when comparing buffer pH (A), oxidant (B), or amine (C). Color indicates the number of mutations in the reads.

Next, we sought to directly compare the performance of two oxidants which have both been previously employed for TimeLapse chemistry. NaIO_4_ is the most common TimeLapse oxidant, but *meta*-chlorobenzoic acid (*m*CPBA) was also characterized to efficiently oxidize s^4^U under TimeLapse conditions (Schofield et al. 2018). In addition, NaIO_4_ oxidizes 3′ diols, requiring the use of *m*CPBA when a 3′ ligation to RNA is required as part of downstream library prep, as is the case in Start-TimeLapse-seq (STL-seq, Zimmer et al. 2021). When comparing TimeLapse-seq data generated with both oxidants, we found that the proportion of total intronic reads containing at least one T-to-C mutation is similar with the two oxidants (Figure 3B). However, *m*CPBA leads to an increase in reads with five or more mutations and a notable increase in the average number of mutations per U (Figures 3B & S3A). We also tested *m*CPBA under near-neutral conditions and found that conversion efficiency was not as high as under acidic conditions (Figures 3B & S3A). Therefore, while *m*CPBA and NaIO_4_ both efficiently convert s^4^U to a cytidine analog, *m*CPBA is slightly more efficient under TimeLapse conditions.

Next, we tested if ammonia could be used as an the amine in TimeLapse-seq as it is expected to convert s^4^U directly to a C instead of a C analog and is used in similar chemistry developed as part of TUC-seq (Riml et al. 2017). Again, we performed NR chemistry on the same RNA using ammonia as the amine, both TimeLapse oxidants, and a basic pH due to the high pKa of ammonia. We found that only NaIO_4_ produced elevated T-to-C mutation rates with ammonia (Figure 3C), showing that the use of a heavy-metal oxidant is unnecessary for nucleotide recoding. When holding the oxidant constant as NaIO_4_, we found that ammonia results in a slightly lower proportion of intronic reads containing a mutation, but the average per U mutation rates are slightly higher than with TFEA as the amine. However, *m*CPBA with TFEA remains the most efficient combination under these conditions (Figures 3B & S3A).

### Estimates of RNA degradation rate constants agree across NR-seq chemistries and are similarly affected by dropout

Finally, we asked if any NR chemistry used in TimeLapse-seq (*m*CPBA + TFEA), SLAM-seq (IAA), and TUC-seq (OsO_4_ + NH_3_) provides a significant benefit and should be preferred in all NR-seq experiments. Previously, TimeLapse-seq, SLAM-seq, and TUC-seq were determined to perform similarly in estimating mRNA degradation rates under standard conditions (Boileau et al. 2021). These experiments were performed using a protocol which minimized experimental droupout by scraping adherent cells from plates prior to lysis; however, the authors did not employ a 3-nt alignment strategy. We used HISAT-3N-aligned data collected with improved handling conditions to compare transcript-specific degradation rate constant estimates (*k*_deg_) obtained with the statistical package bakR between all three NR-seq methods (Vock et al. 2023). In agreement with the previously published comparison, *k*_deg_ estimates produced using data from all three methods strongly agree with each other (Figure 4A-C). We then compared the *k*_deg_ estimates made with data from improved processing strategies to those made with data collected with dropout conditions and aligned with HISAT2 (moderately lenient mismatch penalization). The distributions of the log_2_ fold changes in *k*_deg_ estimates between the two conditions for each NR-seq method is nearly identical, indicating that experimental and computational dropout equally affects *k*_deg_ estimates, independent of NR chemistry (Figure 4D). On average, *k*_deg_ estimates are more than 2-fold higher with optimal conditions than with conditions not optimized to minimize dropout of s^4^U-labeled RNA, demonstrating the importance of minimizing dropout when inferring RNA degradation kinetics.

**Figure 4:**
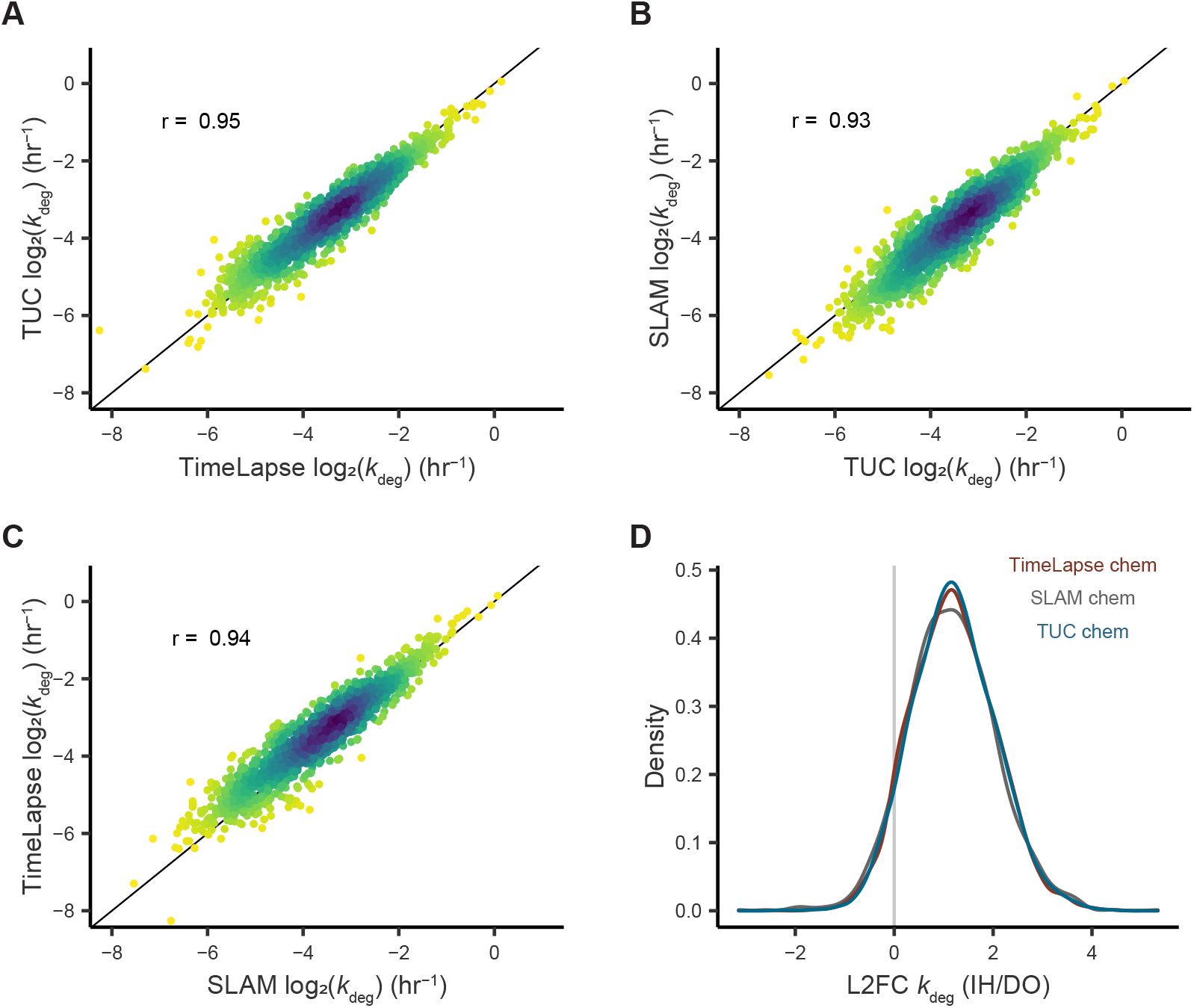
Comparison of RNA degradation rates measured by NR-seq methods and the impact of dropout. (A-C) Scatter plots comparing *k*_deg_ estimates made with bakR using three NR-seq methods. The Pearson correlation coefficient is shown for each. (D) The distribution of the log_2_ fold change in *k*_deg_ estimates when comparing the improved handling and 3-nt alignment (IH) to dropout-vulnerable handling and 4-nt alignment (DO).

## Discussion

We have shown here that analysis of s^4^U-labeled RNA can be biased by RNA sample handling and computational dropout, but that both these sources of dropout can be addressed. s^4^U has long been used as a chemical tool for RNA metabolic labeling, but to our knowledge the specific loss of s^4^U-RNA upon handling RNA samples has not been previously described. This dropout, which happens upstream of NR-seq chemistry, is most likely due to effects such as labeled RNA adhering more strongly than unlabeled RNA to plastic surfaces. This dropout can be avoided with handling precautions such as removing cells from cell culture dishes prior to lysis, and handling labeled RNA in containers with surface treatment designed to minimize the adherence of nucleic acids. Including these simple improvements to protocols can lead to the effective recovery of both s^4^U-labeled and unlabeled RNA from cells. While the focus of this study is on NR-seq approaches, the biochemical dropout described here is upstream of chemical treatments and therefore is likely to impact any experiment that relies on comparisons between s^4^U-labeled and unlabeled RNA. In all such experiments, it is important to sequence samples with and without the metabolic label to control for any potential biological impact of the label as well as the dropout described here.

Furthermore, T-to-C mutation-containing reads in sequencing data are more difficult to align due to mismatch penalties applied by standard RNA-seq aligner software. We showed that reads with more T-to-C mutations are less likely to be aligned with a standard RNA-seq aligner. This is at least partially attributable to standard aligners requiring a seed sequence to perfectly match the reference genome (Berg et al. 2023). Typically the seed is twenty base pairs long, but if a perfect twenty base pair long stretch does not exist, the read will fail to be aligned. In addition, each mismatch between the read and reference genome incurs an alignment penalty. If there are a sufficient number of mismatches, the read may fail to align. Each of these parameters can be customized to a certain extent to allow for more mismatches, but this also allows for mismatches of any N-to-N and not just T-to-C. To solve this issue, we took advantage of a new 3-nt RNAseq aligner, HISAT-3N, which does not penalize the induced T-to-C mismatches (Zhang et al. 2021). HISAT-3N follows a similar strategy as aligners developed for bisulfite sequencing: all Ts in the reference genome and sequencing data are converted to C’s. The 3-nt alignment prevents T-to-C mismatches from penalizing alignments, and because HISAT-3N stores information about the original sequence, it can be used to identify T-to-C mutations. HISAT-3N was previously demonstrated to be a fast and accurate aligner, and is therefore highly recommended for all NR-seq experiments requiring a splice-aware aligner.

RNA molecules with short average half-lives will be more highly labeled with s^4^U than those with long average half-lives. Consequently, each fast-turnover RNA molecule is more likely to be lost during handling, and reads derived from these molecules are more likely to misalign during computational processing. Therefore, dropout introduces a bias against transcripts with short half-lives, an effect that we found to be prevalent within published datasets. Ultimately, dropout leads to an overestimation of transcript half-lives and a dampening of any changes in turnover rates caused by an experimental treatment. Fortunately, we were able to show that our improved handling and computation protocols fully reversed this bias such that NR-seq data was essentially indistinguishable from traditional RNA-seq data in terms of read counts and intronic content.

We showed that TimeLapse chemistry efficiently converts s^4^U to a cytidine analog with either NaIO_4_ or *m*CPBA and performed best under slightly acidic conditions. *m*CPBA achieved a modestly higher conversion rate than NaIO_4_ and preserves 3′ ends of RNAs, which can be important for downstream processing steps of a sequencing experiment (Zimmer et al. 2021). Therefore, the difference between NaIO_4_ and *m*CPBA is inconsequential unless a 3′ adapter ligation is required. Likewise, TFEA and ammonia can be used as the nucleophilic amine with NaIO_4_, but ammonia should not be used with *m*CPBA.

Finally, s^4^U recoding chemistries developed as part of TUC-seq, SLAM-seq, and TimeLapse-seq tend to provide comparable estimates of the steady-state kinetics of cellular RNA, unless material availability or safety is a concern. Independent of chemistry selection, additional consideration must be taken when preparing samples and aligning NR-seq data. Without awareness of dropout, high levels of dropout caused by unoptimized RNA sample handling could easily be misinterpreted as biological effects of labeling. NR-seq is a quickly evolving technique that has been extended to numerous applications such as analysis of the kinetics of sub-cellular RNA trafficking (Smalec et al. 2022), and methods to analyze NR-seq data are still being developed (Qiu et al. 2022; Vock et al. 2023). With the work presented here, we establish new guidelines to address previously unappreciated challenges of performing NR-seq which will improve the power of NR-seq methods as tools to study RNA dynamics.

## Materials and Methods

### Cell lines and s^4^U metabolic labeling

Metabolic labeling of cells was performed as described previously (Schofield et al. 2018). For all NR-seq experiments, HEK293T cells were grown to approximately 70% confluency when the media was spiked with s^4^U (100 μM). For NR-seq experiments with improved handling conditions, plates were immediately placed on ice and washed with ice-cold PBS. Cells were scraped from plates, transferred to low nucleotide-binding tubes, and pelleted by centrifuging at 500Xg for 5 min. PBS was removed and cells were resuspended in 1 mL TRIzol and frozen at -80°C. For all NR-seq experiments with dropout handling conditions, ice cold TRIzol was added directly to plates, pipetted up and down to spread across the plate, and left on ice for 5 min to fully lyse the cells. Cellular lysate was then transferred to a separate tube and frozen at -80°C.

### NR-seq (TimeLapse-seq, SLAM-seq, TUC-seq)

Genomic DNA was depleted by treating with TURBO DNase and total RNA was extracted with one equivalent volume of Agencourt RNAClean XP beads according to manufacturer’s instructions. 5 μg of total RNA was subjected to TimeLapse, SLAM, or TUC chemistry as previously described with some modifications (Schofield et al. 2018; Herzog et al. 2017; Riml et al. 2017).

For TimeLapse-seq, RNA was mixed with 600 mM TFEA or NH_3_, 1 mM EDTA and 100 mM sodium acetate pH 5.2 or 100 mM Tris pH 7.4. Then, NaIO_4_ or *m*CPBA was added to 10 mM final and the reaction was incubated at 45°C for 1 h. RNA was purified with one volume of Agencourt RNAClean XP beads and eluted with nuclease-free water. RNA was mixed with 10 mM DTT, 10 mM Tris pH 7.4, 5 mM EDTA, and 50 mM NaCl and incubated at 37°C for 30 min. RNA was purified with one volume of Agencourt RNAClean XP beads and eluted with nuclease-free water. For SLAM-seq, RNA was mixed with 50% DMSO, 50 mM sodium phosphate buffer pH 8.0 and 10 mM IAA and incubated at 50°C for 15 min. The reaction was stopped by adding excess DTT. RNA was purified with one volume of Agencourt RNAClean XP beads and eluted with nuclease-free water.

For TUC-seq, RNA was mixed with 180 mM NH_4_Cl and 450 μM OsO_4_ and incubated at 25°C for 3 h. RNA was purified with one volume of Agencourt RNAClean XP beads and eluted with nuclease-free water.

For each sample, 10 ng of RNA input was used to prepare sequencing libraries from the Clontech SMARTer Stranded Total RNA-Seq kit (Pico Input) with ribosomal cDNA depletion. Libraries were sequenced on a NovaSeq 6000 2X100bp.

### NR-seq alignment and mutation calling

Filtering and alignment to the human GRCh38 genome version 26 (Ensembl 88) or the *Drosophila* dm6 genome (or a combined genome when using spike ins for normalization) were performed as described above for STL-seq with some differences. Reads were trimmed of adaptor sequences with Cutadapt v1.16 (Martin 2011) and aligned to GRCh38 or dm6 using HISAT-3N (Zhang et al. 2021), HISAT2 (Kim et al. 2019), or STAR (Dobin et al. 2013). For HISAT-3N, default parameters and --base-change T,C were used. Parameter settings used for HISAT2 and STAR are described in Figure 2A. Reads aligning to transcripts were quantified with HTSeq (Anders et al. 2015) htseq-count. SAMtools v1.5 (Li et al. 2009) was used to collect only read pairs with a mapping quality greater than 2 and concordant alignment (sam FLAG = 99/147 or 83/163). Mutation calling was performed essentially as described previously (Schofield et al. 2018). Briefly, T-to-C mutations were only considered if they met several conditions. Mutations must have a base quality score greater than 40 and be more than 3 nucleotides from the read’s end. Sites of likely single-nucleotide polymorphisms (SNPs) and alignment artifacts were identified with bcftools or from sites of high mutation levels in the non-s^4^U treated controls (binomial likelihood of observation p ¡0.05). These sites were not considered in mutation calling. Browser tracks were made using STAR v2.5.3a (Dobin et al. 2013). Normalization scale factors were calculated with edgeR (Robinson et al. 2010) using read counts from the spike-in species (calcNormFactors using method = ‘upperquartile’). If using spike ins for normalization, only reads aligning to the genome of the spike in species were used for normalization with edgeR.

### Estimation of RNA decay and synthesis kinetics

For all kinetic analyses of NR-seq data, the bakR R package was used to estimate RNA degradation rates and the observed changes in these rates upon changes in sample and data handling (Vock et al. 2023). Default settings were used with the MCMC model (StanFit = TRUE). RNA synthesis rates and changes in synthesis rates were determined as outlined in the bakR manual using DESeq2 to estimate changes in total RNA.

## Data and code availability

Data sourced from published available literature are available on Gene Expression Omnibus (GEO) under GSE162264 (Narain et al. 2021), GSE200422 (Fisher et al. 2022), GSE156885 (Richters et al. 2021), or GSE139151 (Zuckerman et al. 2020). Raw and processed NR-seq data generated in this study are available and accessible on GEO under GSE233139. All NR-seq data were analyzed with the TimeLapse pipeline v0.4 (TimeLapse pipeline v0.4, bitbucket.org/mattsimon9/timelapse_pipeline/src/master/) and the bakR R package (https://github.com/simonlabcode/bakR/) which are both publicly available. In addition, to facilitate preprocessing of NR-seq data, we recently released a Snakemake (Mölder et al. 2021) implementation of our alignment and mutation counting pipeline which is also publicly available (https://github.com/simonlabcode/bam2bakR).

## Acknowledgments

We thank K. Neugebauer, J. Steitz, S. Slavoff, N. Dimitrova, and all of the members of the Simon Lab for valuable discussions and feedback. This work was supported by the NIH NIGMS T32GM007223 (to J.T.Z., I.W.V., and J.A.S), an AHA Predoctoral Fellowship (to L.K.), and NIH R01 GM137117 (to M.D.S.).

## Author contributions

Conceptualization: J.T.Z., I.W.V., J.A.S., L.K., M.H.M., M.D.S.; Methodology: J.T.Z., J.A.S., L.K., M.D.S.; investigation: J.T.Z., I.W.V., J.A.S., L.K.; formal analysis: J.T.Z., I.W.V.; resources: M.D.S; validation: J.T.Z., I.W.V., J.A.S., L.K., M.H.M.; writing: J.T.Z., I.W.V., M.H.M., M.D.S.; visualization: J.T.Z., I.W.V., M.D.S.; supervision: M.D.S.; project administration: M.D.S.; funding acquisition: M.D.S.

---

**Figure S1:**
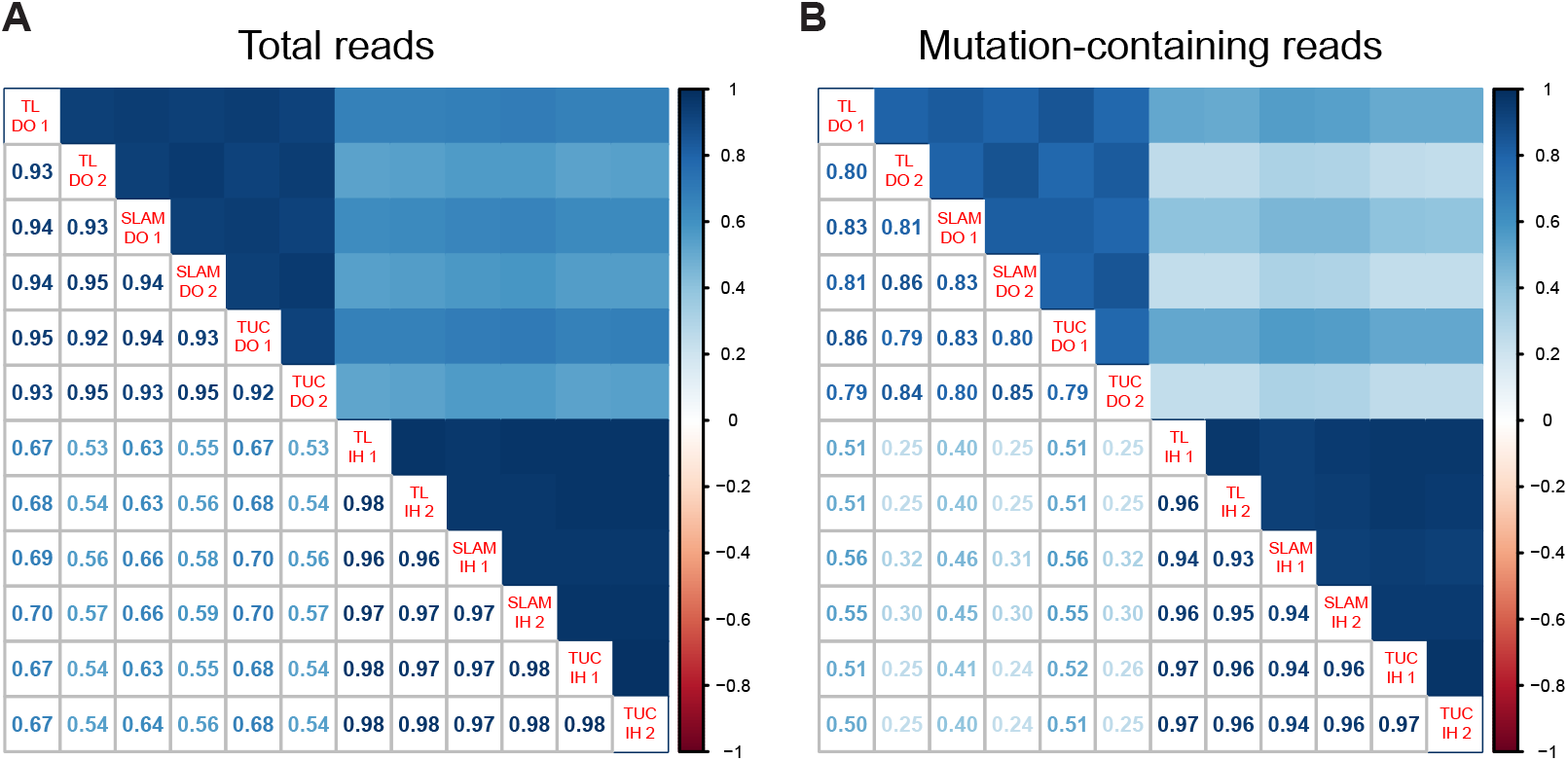
NR-seq data prepared with different handling conditions do not correlate well with each other. (A,B) Correlation plot of the log_2_ counts of all (A) or T-to-C mutation containing (B) reads aligning to genes in NR-seq data with both dropout (DO) and improved handling (IH) conditions and aligned with HISAT2.

**Figure S2:**
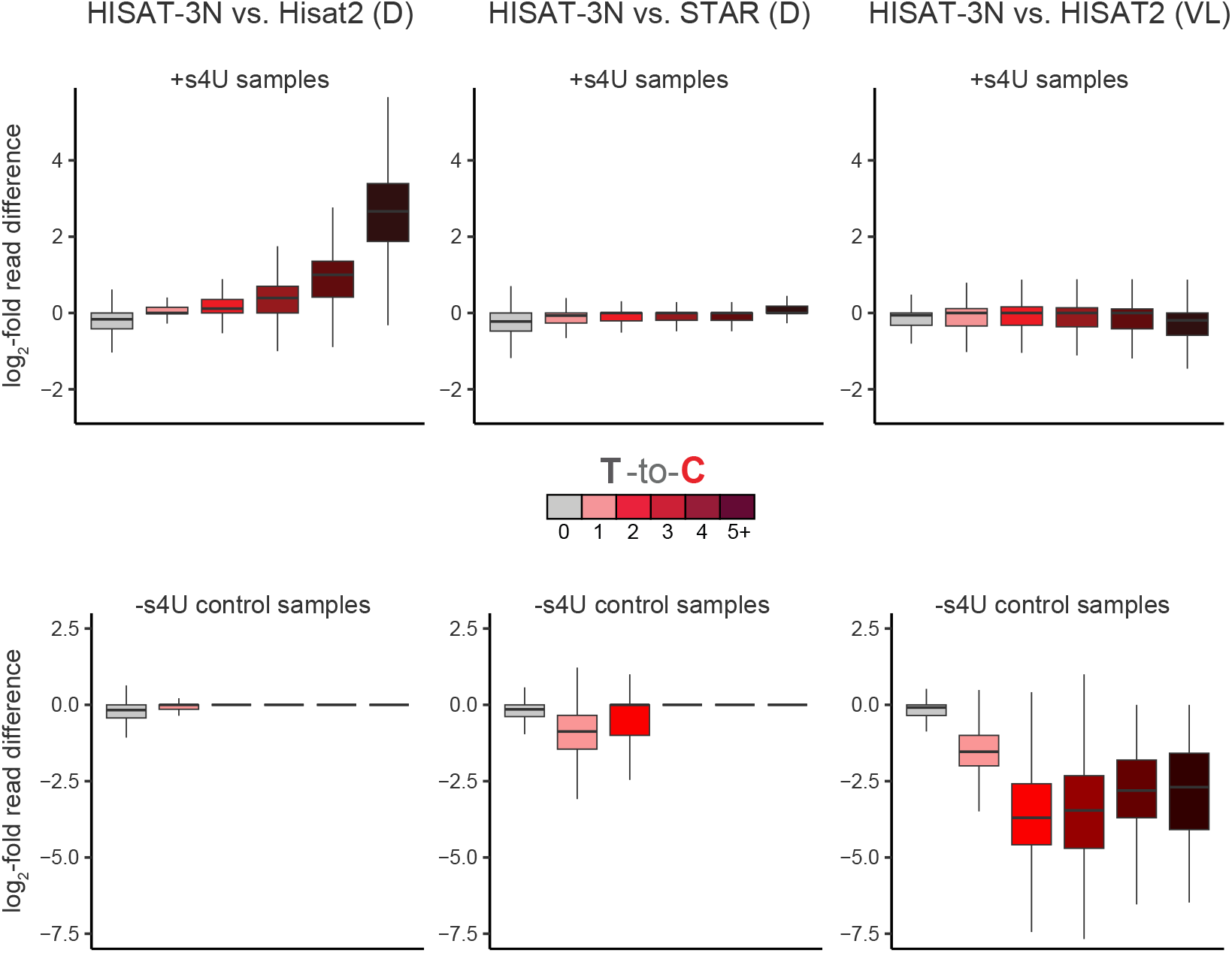
3-nt alignment increases mutational sensitivity while not sacrificing specificity. Boxplots of transcript-specific differences in the number of reads with a particular number of T-to-C mismatches. Left column compares HISAT-3N to HISAT2 with default settings. Middle column compares HISAT-3N to STAR with default settings. Right column compares HISAT-3N to HISAT2 with extreme mismatch leniency. Top row is data for s^4^U fed samples. Bottom row is for unfed control samples. Y-axes are log_2_-fold differences between HISAT-3N and the alternative aligner in the number of reads with a T-to-C mismatch count indicated in the color bar.

**Figure S3:**
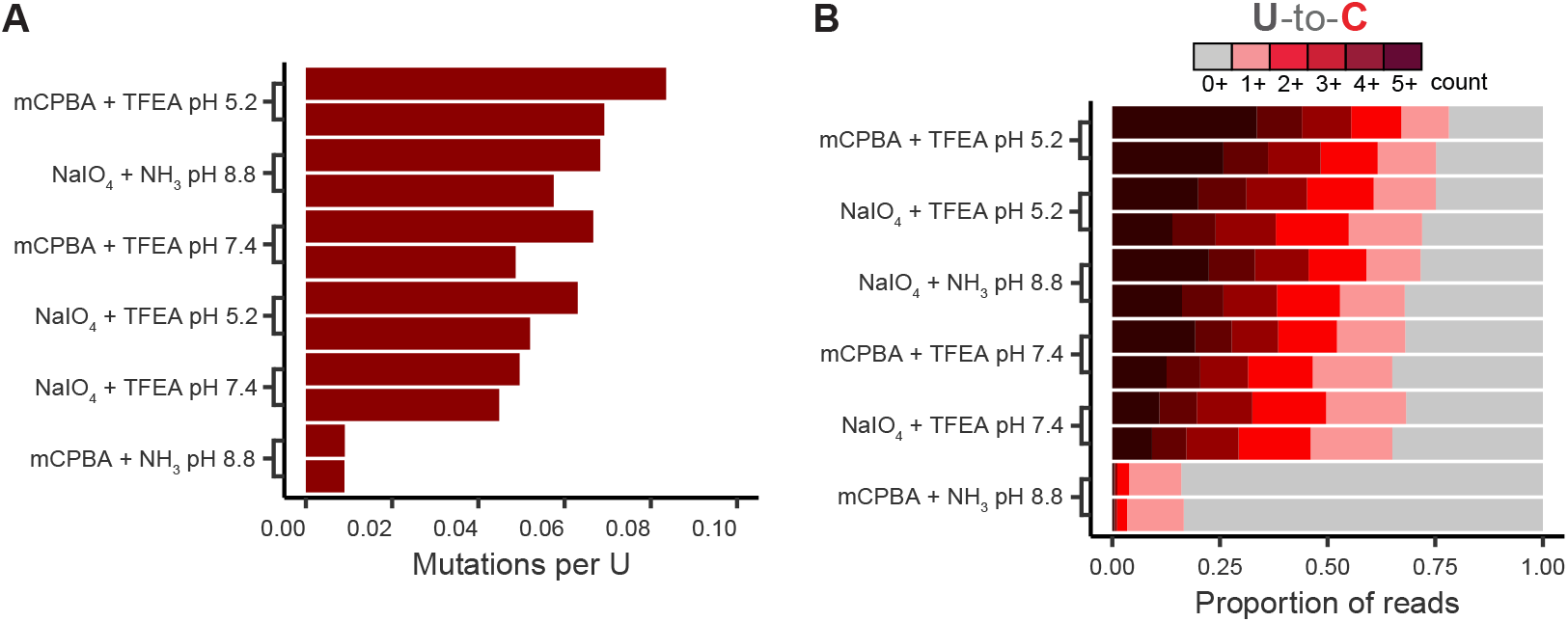
Comparison of mutational content in TimeLapse-seq data using different reaction conditions. (A) The per U mutation rate in all intron-aligning reads with different TimeLapse conditions (B) The proportion of intron-aligning reads which contain T-to-C mutations with different TimeLapse conditions. Color indicates the number of mutations in the reads.

